# Improving RNA Secondary Structure Prediction Through Expanded Training Data

**DOI:** 10.1101/2025.05.03.652028

**Authors:** Conner J. Langeberg, Taehan Kim, Roma Nagle, Charlotte Meredith, Dimple Amitha Garuadapuri, Jennifer A. Doudna, Jamie H. D. Cate

## Abstract

In recent years, deep learning has revolutionized protein structure prediction, achieving remarkable speed and accuracy. RNA structure prediction, however, has lagged behind. Although several methods have shown some success in predicting RNA secondary and tertiary structures, none have reached the accuracy observed with contemporary protein models. The lack of success of these RNA structure prediction models has been proposed to be due to limited high-quality structural information that can be used as training data. To probe this proposed limitation, we developed a large and diverse dataset comprising paired RNA sequences and their corresponding secondary structures. We assess the utility of this enhanced dataset by retraining on a deep learning model, SincFold. We find that SincFold exhibited improved generalization to some previously unseen RNA families, enhancing its capability to predict accurate de novo RNA secondary structures. The RNASSTR dataset provides a substantial advance for RNA structure modeling, laying a strong foundation for the development of future RNA secondary structure prediction algorithms.

## Introduction

Structured RNAs play essential regulatory roles across all domains of life and in viruses (1, 2), and participate in diverse regulatory processes, including transcription and translation (2, 3), catalysis (4–6), epigenetic modulation (7), and ribonucleoprotein complex function (8, 9). Similarly, many viruses make use of structured RNA motifs during infection to enhance virulence and replication efficiency (10, 11). Recent advances in mRNA vaccines have specifically leveraged RNA structural stability to enhance half-life and protein expression (12), highlighting the role of RNA structure in molecular therapeutics. The utility of RNA structure prediction represents a promising frontier for antiviral drug design (13, 14), RNA-targeting small molecules (13), CRISPR guide RNA design (15, 16), and RNA-based synthetic biology applications (17–19). However, the effective use of RNA in these areas necessitates a robust and comprehensive understanding of RNA folding and structure.

Structure determination methods including X-ray crystallography, cryo-electron microscopy, and nuclear magnetic resonance spectroscopy (20–25) are the only methods which allow direct experimental confirmation of an RNA’s structure, including the three dimensional topology and interaction network within a fold. These methods are essential in defining the complex, non-Watson Crick Franklin (WCF) type base pairs, long-range interactions, and pseudoknots which define the higher order topology of structured RNA. However, these methods are resource-intensive and frequently fail due to technical limitations or inadequate resolution for directly modeling the RNA structure. High-throughput methods such as DMS-MaP Seq (26, 27) and SHAPE-MaP (28–32) allow for efficient scaling of experiments but sacrifice atomic-level resolution, relying heavily on expectation-maximization algorithms that may obscure non-standard structural features.

Inspired by the recent success of deep learning methods for protein structure prediction like AlphaFold2 (33) and ESMFold (34), efforts have been directed at applying similar approaches to RNA structure prediction (35). However, the limited availability of experimental RNA structures significantly hampers these data-intensive approaches, making it challenging to achieve comparable accuracy to protein predictions (36, 37). Beyond the challenges posed by the limited training data for RNA 3D structure prediction, the multiple sequence alignment algorithms used for protein structure prediction do not work well for RNA (38, 39). Protein sequence alignments leverage evolutionary conservation that can be detected in the primary structure directly, further informing structural conservation and relationship with other homologs.

RNA alignments however, are dominated by secondary structure due to the coevolution of paired bases (39, 40) which appear as compensatory mutations. While compensatory mutations conserve structure, they may result in highly degenerate sequences that are therefore difficult to align. RNA relies on base pairing to form the initial topology of the fold, necessitating secondary structure informed sequence alignments rather than relying on the primary structure. Accurate RNA secondary structure prediction is therefore an essential prerequisite to accurate 3D structure prediction as the WCF base pairs define the stems, junctions, and pseudoknots in the RNA structure around which the tertiary contacts form (41, 42).

Historically, a combination of experimental and bioinformatic methods have been used to infer RNA secondary structure (39, 43–47). However, these approaches largely require specific expertise, making them hard to disseminate and to scale. Presently-available computational algorithms that aim to minimize the requirement for user expertise while providing accurate predictions frequently fail to accurately predict large, multistem RNA folds due to their reliance on reductionist thermodynamic models which do not recapitulate the often many and non-nested stems present in highly structured RNA. For example, computational methods such as ViennaRNA (RNAfold) and MFold (48, 49), rely heavily on the Turner Rules (46, 50–54), utilizing empirically determined nearest-neighbor thermodynamic parameters to minimize the folding free energy. Although these programs proved effective for motifs with fewer stems and shorter lengths, they often perform poorly on long RNAs containing many stems and pseudoknots due to the oversimplified energetic approximations and folding assumptions. Recent machine learning methods, including UFold, SincFold, and DMfold (55–57), integrate deep neural networks to enhance the accuracy of RNA secondary structure prediction. However, these deep learning models frequently exhibit poor generalization (42), performing well only on RNA folds the models are directly trained on. This can be observed in careful validation where existing models demonstrate a significantly decreased prediction accuracy with RNA folds not used in model training (42). More recently, hybrid methods such as the MXFold suite and CDPfold (58, 59), which integrate both machine learning and thermodynamic approaches have shown promise in improving model performance. However these methods have not yet achieved accurate and general RNA secondary structure prediction (38, 42). Integrating supplementary data has been suggested as a method to improve model performance, such as the use of chemical probing or enzymatic mapping data (14, 60). The scalability of these methods enabled by high throughput sequencing and ease of data generation make integrating these orthogonal approaches a promising future direction in RNA structure prediction.

One possible solution to improve present machine learning models for RNA secondary structure prediction is to increase the size and diversity of training data. The existing training data, such as bpRNA (61) and ArchiveII (41) contain a limited set of distinct RNA folds, few sequences, and highly biased compositions which may encourage machine learning models to memorize the most abundant classes, limiting their usability with novel RNA folds and motifs. ArchiveII, for example, consists of 10 distinct RNA folds and 3847 sequences whereas the version 14.10 of the Rfam (62, 63) database recognises 4170 distinct RNA families. The bpRNA dataset provides a more structurally diverse set of sequence structure pairs than ArchiveII, with 2,588 structural families from Rfam, though only consisting of 102,318 sequences. Additionally, the data lack a standardized grammar for model training, which makes comparisons between users challenging to achieve. Finally, there is a need for robust methods to prepare training data that accounts for both sequence and structural diversity in splitting RNAs into the training and test data.

To overcome these challenges, we have developed the RNA Secondary Structure Repository (RNASSTR), a rigorously curated dataset comprising 4170 annotated RNA families described in Rfam and nearly 5 million unique sequences. These sequences span all domains of life and viruses, and were assembled leveraging robust bioinformatic workflows to identify novel RNA structural homologs. Using RNASSTR, we retrained two existing RNA secondary structure prediction models, demonstrating that increased depth and diversity of RNAs may improve model generalization, but at the cost of performance for some structural families. Our analysis also identifies additional limitations suffered by current models including slow training speed and sequence-structure memorization. The RNASSTR dataset and associated benchmarking model parameters thus provide a powerful foundational resource that can be used for further model development, and represents an important step in developing training data comparable to those now available for proteins.

## Materials and Methods

### Dataset Generation

To generate sequence-structure pairs we leveraged the bioinformatic tool Infernal v1.1.5 and the RNA Family database (Rfam) v14.10 (47, 63). Using the 4170 covariance models of RNA structure families deposited on Rfam, we searched against a database containing all eukaryotic and viral reference genomes from NCBI, release 229 (64), as well as all bacterial and archaeal reference genomes from the Genome Taxonomy Database, release 214 (65). The resulting sequences were realigned to their respective Rfam families using Infernal, thus inferring secondary structure from the consensus model. The resulting hits were then filtered by several features. First, only sequences with a reported E-value of 0.01 or less were considered, an approximate false positive rate of 1 out of 100 or better. Sequences more than 2 standard deviations from the Rfam defined length were removed, in line with previously reported filtering thresholds (66). Similarly, sequences with more than 2 standard deviations below Rfam defined consensus base pairs were removed as well as those sequences with more than 2 standard deviations fewer WCF base pairs than defined in the Rfam seed alignment were removed. Sequences which were identified outside of their Rfam defined phylogeny were removed. Finally, overlapping hits were assessed and only the better E-value sequence was retained.

In order to facilitate rigorous training and prevent data leakage we performed data splitting accounting for secondary structure. To do this we defined mutually exclusive RNA structural groups using Infernal cross-validation, identifying RNA families incapable of cross-identification, thus preventing structural data leakage. This was accomplished by searching all sequences of one Rfam family against all other Rfam models. If any sequences were identified as having statistically significant similarity, an E-value less than or equal to 0.01, they were considered not mutually exclusive and were placed into one of the data splits. Those families which did not have identifiable structural homology were placed in different data splits. This resulted in a partitioning scheme containing approximately 90% training, 5% validation, and 5% test sequences, with exact counts provided in the supplementary materials. For model training purposes, sequence-structure pairs were converted into several formats using a custom python script: standard FASTA (67) and dot-bracket notation, BPSEQ format (61), and an expanded BPSEQ format specific to the SincFold (56) we here call SincFold format.

### SincFold retraining

Model retraining was performed on a single NVIDIA RTX A4500 GPU. Default parameters were used for model training from a random seed initialization, as specified in the initial publication (56). This ensured models were trained from scratch without any prior knowledge. Training was monitored using the calculated F1 score of the train and validation split at the end of each epoch of training to externally monitor model progress. Training was allowed to progress until the validation F1 score plateaued for multiple epochs, in this case training required 15 epochs to stabilize as we observed early convergence. The final trained model was assessed using the calculated F1 score of the train and test split.

### MFold secondary structure calculation

MFold v3.6 (49) was used to calculate the minimum free energy structures of a subset of each RNA family to compare against the ML models. A subset of 100 sequences was randomly selected from each RNA family and subjected to folding using the default parameters in mFold. For those RNA families with less than 100 members all sequences were used.

### F1 and MCC score calculation

Both F1 and Matthews Correlation Coefficient (MCC) scores were calculated using a custom script which allowed us to analyze all predicted secondary structures independently. In order to ensure a rigorous calculation, base pair partners were enumerated using the same strategy the BPSEQ format uses. From these enumerated pairing schemes, both the F1 and MCC scores were calculated. A true positive is defined as a nucleotide predicted to be involved in a base pair with the correct pairing partner, a false positive is defined as a nucleotide predicted to be involved in a base pair but with the incorrect pairing partner, and a false negative is the number of base pairs in the ground truth structure not predicted. True negatives are ignored for this adaptation of MCC as is standard practice in RNA 2D structure quantification.

True Positive (TP): Predicted base pair is in the true structure.

False Positive (FP): Predicted base pair is not in the true structure.

False Negative (FN): True base pair was not predicted.

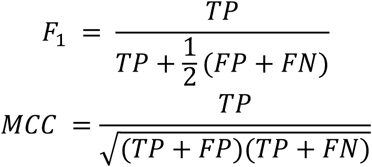

### Models which could not be retrained

In addition to retraining both SincFold, we attempted to retrain a number of other ML models. However, these were unable to be retrained for various reasons ranging from missing training scripts to bugs in the deposited code. The following is an overview of those models which we attempted to retrain and why we were unable to do so.

- UFold (55): During model training, an error arose stating the input dimension did not match the shape of the data preventing the model from being retrained. This error could not be resolved. GitHub link: https://github.com/uci-cbcl/UFold
- REDfold (68): Script typos prevented the retraining script from functioning. GitHub link: https://github.com/aky3100/REDfold
- SPOT-RNA (69): Model lacked a retraining script. GitHub link: https://github.com/jaswindersingh2/SPOT-RNA
- E2Efold (70): Unresolvable munch incompatibility prevented model retraining. GitHub link: https://github.com/ml4bio/e2efold
- MXFold2 (58): Training times were prohibitively long, approaching 270 hours per epoch. GitHub link: https://github.com/mxfold/mxfold2

## Results

### Dataset curation

In order to construct a deeper and more diverse RNA secondary structure dataset compared to those presently available, we leveraged the existing bioinformatic tool Infernal to gather and curate a set of structurally homologous RNA sequences. We first retrieved all covariation models from Rfam version 14.10 (63), which is composed of 4170 covariation models of structured RNAs (39). These models describe the sequence and secondary structure space of each unique RNA family using a stochastic context free grammar and can be used by the bioinformatic tool Infernal (47) to search for sequences which can adopt a similar secondary structure topology. We then used these covariation models to search all reference genomes available from the GTDB (version 214) (65) and NCBI RefSeq databases (resease 229) (64), from which we identified 8,910,328 putative homologous sequences (Fig. 1A,B). Because of the underlying false positive rate inherent in these search strategies, we chose to refine the dataset using a number of metrics to minimize the inclusion of false positive structures in the final dataset. We curated this set of data using a number of statistical metrics as described in the methods section, based on a similar analysis used for a prior dataset (66) (Fig. 1C). Briefly, we removed those sequences which we determined to be outliers in terms of sequence length and structure conservation, as well as those which fell outside of the expected phylogeny as defined by Rfam. Following this curation process, we recovered 4,779,435 high confidence RNA sequence-structure pairs. At present, the resulting dataset, RNASSTR, does not include pseudoknots due to the limitation that the stochastic context free grammars used by Infernal can only handle nested stems and as such cannot be used to identify pseudoknots (47).

**Figure 1.**
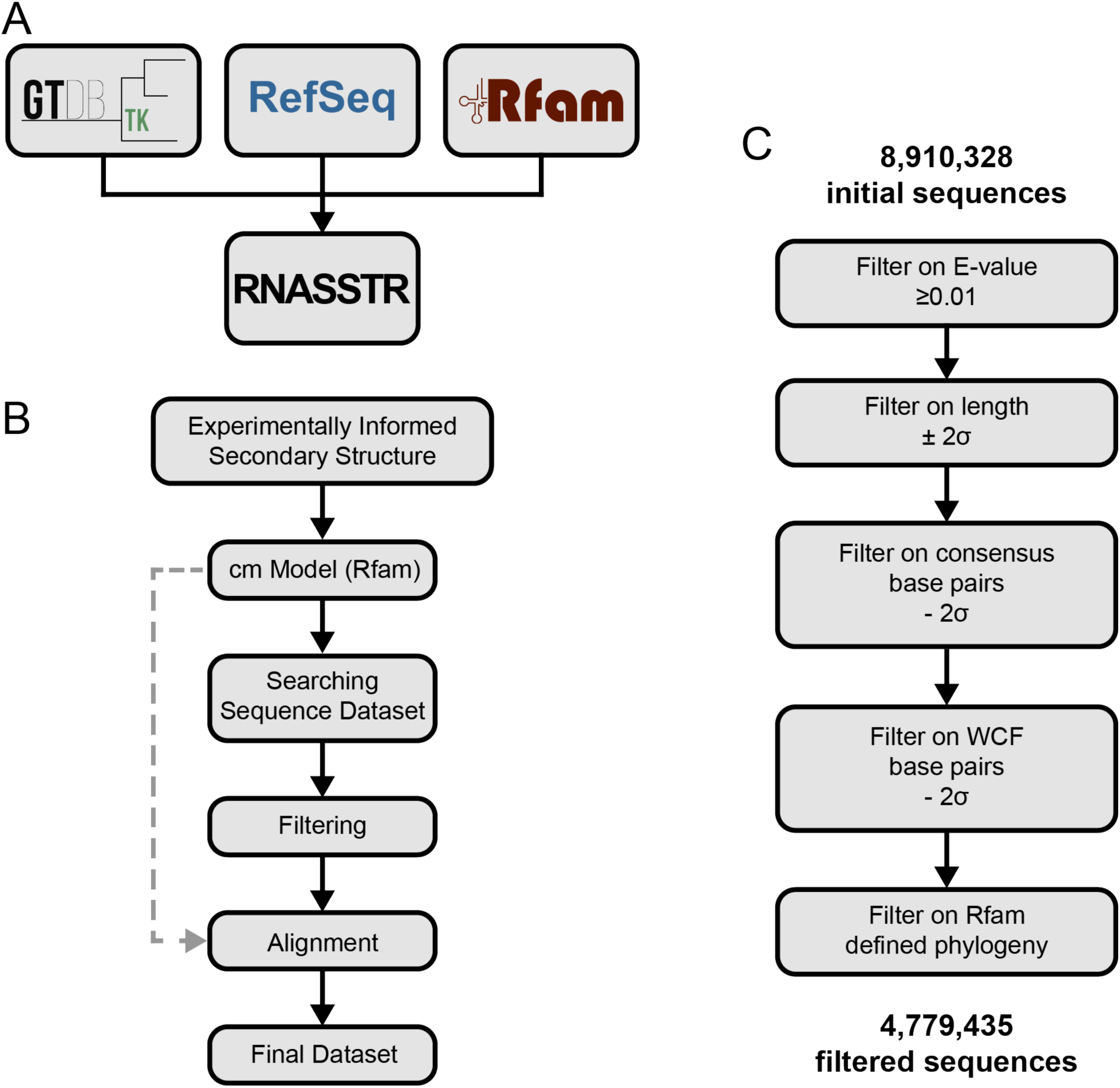
RNASSTR dataset generation and parsing. **A)** RNASSTR contains reference genomes from both the Genome Taxonomy Database (GTDB) and NCBI RefSeq datasets, together consisting of reference genomes from all domains of life as well as viruses. The RNA family database, Rfam, contains covariation models of 4170 RNA structure families. **B)** The stepwise bioinformatic workflow used to generate RNASSTR including search, parsing, and data splitting. **C)** The step-by-step process for filtering putative RNA structure homologs for RNASSTR dataset generation.

Further analysis of the resulting dataset revealed that the bulk of the sequences were centered around 80 nucleotides in length, composed largely of tRNAs, pre-miRNAs, and bacterial sRNAs (Fig. S1). In addition to this large population, several other notable populations exist, such as the bacterial small subunit ribosomal RNA at approximately 1600 nucleotides in length. Notably, the composition of RNASSTR is not equally distributed between the distinct RNA families. A subset of these families constitutes the bulk of the sequences, with the tRNA family accounting for 39.5% of sequences within the dataset (Fig. 2A,B). However, when compared to other frequently used RNA sequence structure training datasets RNASSTR provides a greatly increased depth across other RNA families for model training (Fig. 2C). In addition to the significant increase in sequence-structure pairs over other datasets, RNASSTR also demonstrates superior sequence diversity at a variety of fractional sequence identities, as can be seen in the top 6 most abundant RNA families from RNASSTR (Fig. 2D). Because the goal of these RNA secondary structure prediction models is to accurately predict the native fold of a given RNA, ideally generalizing to unseen structural classes, we partitioned the dataset into three parts, a training set containing 90% of sequences belonging to one-third of the Rfam families, a validation set containing 5% of the sequences belonging to one-third of the Rfam families, and a test set containing the final 5% of sequences and one-third of Rfam families. By ensuring that the families were mutually exclusive at the structural level we ensured no data leakage between the training, validation, and testing sets, which should aid model generalization and mitigate against memorization.

**Figure 2.**
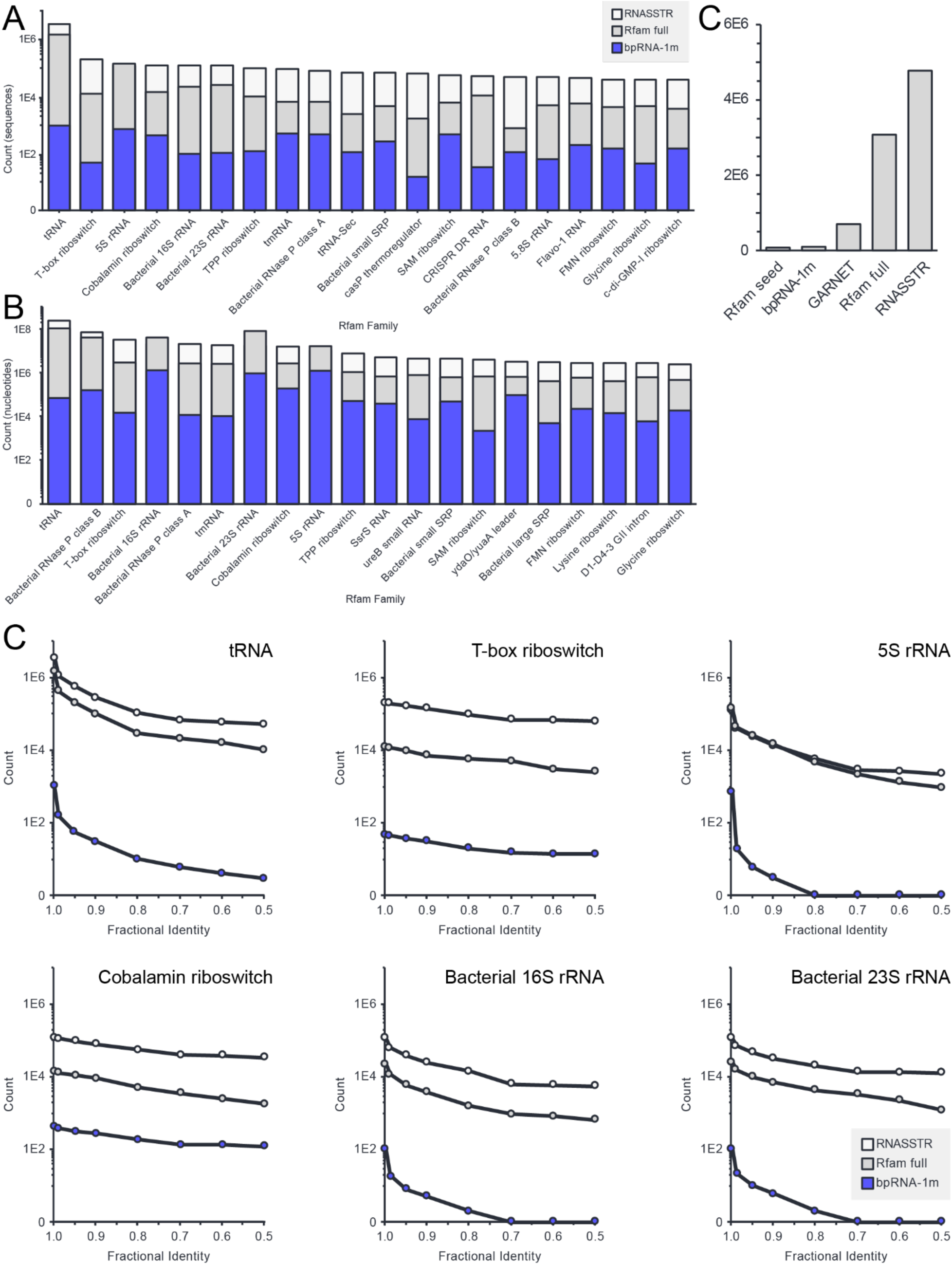
Depth and diversity of RNASSTR RNA sequence-structure pairs. **A)** Stacked histogram of the top 20 most abundant classes in RNASSTR by sequences compared to Rfam full and bpRNA-1m. **B)** Stacked histogram of the top 20 most abundant classes in RNASSTR by nucleotides compared to Rfam full and bpRNA-1m. **C)** Graphical representation of the total abundance of sequence-structure pairs in multiple RNA secondary structure datasets. **D)** Fractional identity of six abundant classes of RNA families comparing the sequence depth at multiple thresholds comparing RNASSTR, Rfam full, and bpRNA-1m.

### Model retraining

We next tested whether using RNASSTR to retrain existing machine learning algorithms would improve their performance and generalization. To do this we identified a previously published 2D RNA prediction model, SincFold (56), which we could retrain from scratch. We used the structure-stratified dataset split within RNASSTR to prevent data leakage between training and testing (Methods). We trained SincFold until the validation F1 score converged, to a training F1 of 0.983 and a validation F1 of 0.420, which took 15 epochs. Notably, one full training and validation cycle for SincFold required 64 GPU hours per training epoch and 2 hours per validation iteration.

### Model performance

Following model training, we used the retrained SincFold model as well as its default published parameter sets to predict the secondary structures of the RNASSTR held-out testing data. We measured the performance using two standard metrics for the field, F1 and MCC scores, both measures of confusion matrix categories (Methods). We computed these metrics for the published model parameters and for our retrained parameter sets to assess how data scaling during training affected the model’s ability to generalize to unseen RNA families and folds. To compare against non-ML based methods, we included predictions from a popular minimum free energy program, RNAfold (49). For RNAfold, we subsampled 100 sequences per family in the testing set due to the computational complexity of computing all structures.

For SincFold, the retrained model performed worse than the published model on the RNASSTR testing set. This is striking, given that we made no changes to the underlying model architectures or hyperparameters. The retrained SincFold model did, globally, outperform RNAfold, the minimum free energy model (Fig. 3A,B, Table 1). To better understand if specific features of the dataset impacted the retrained SincFold model performance, we assessed a range of sequence features: absolute number of ground truth paired bases, GC content, ground truth fraction of paired bases, and sequence length. While no clear trend is visible in the data, we noted many sequences with a F1 score of 0 indicating the retrained SincFold model failed to predict any true positives (Fig. S3), defined as correctly predicted base pairs. In these cases, the retrained model appears to catastrophically fail during the inference stage.

**Figure 3.**
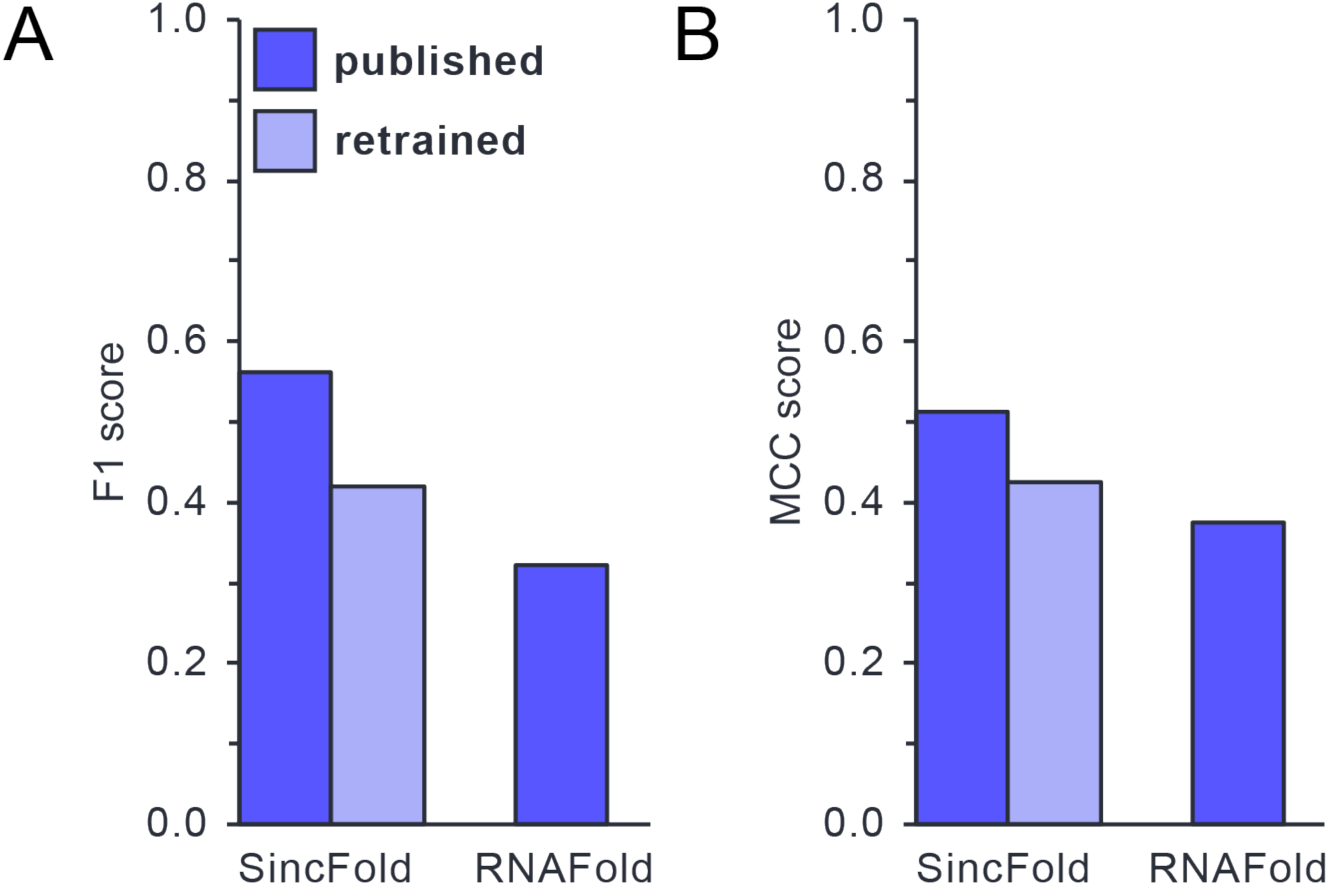
Model performance pre and post retraining. **A-B)** Model performance on RNASSTR test partition using the published model and RNASSTR retrained model for SincFold. RNAfold is included as a minimum free energy comparison. Scores are calculated for F1 (**A**) and MCC (**B**).

**Table 1.** Model performance. Shown are the RNASSTR test partition for published and RNASSTR models as well as minimum free energy model RNAFold.

|  | SincFold<br>published | SincFold<br>RNASSTR<br>retrained | RNAFold |
| --- | --- | --- | --- |
| Test F1 | 0.561 | 0.420 | 0.321 |
| Test MCC | 0.513 | 0.424 | 0.391 |

We then assessed per-family F1 scores to determine if specific RNA structures were more prone to poor prediction accuracy. We observe that the distribution of F1 scores for the top 20 families did not change significantly across epochs (Fig. S4). However, in later epochs the retrained SincFold model showed improvement in performance for some Rfam groups, RF01359 and RF00730 as examples, suggesting that the retrained SincFold model learned to better predict some Rfam groups beyond early epochs. While this was true for a subset of classes, most Rfam groups are already learned at early epochs.

When comparing the published SincFold model to the RNASSTR retrained model, we observed a large variance in per family performance (Fig. 4A,B), suggesting these two models learned different features resulting in varying results. While the published model parameters yielded a marginally higher average F1 score across the full test set, this improvement was spread thinly across many RNA families. In contrast, the RNASSTR retrained model showed more substantial gains within a smaller subset of families, suggesting that although it performs less consistently overall, it captures specific structural features more effectively. This implies that while the retrained model does not perform as well broadly, within specific classes the retrained model performs better. To exemplify this we show two representative sequences, showing the ground truth structure, the published SincFold prediction, and the RNASSTR retrained SincFold prediction (Fig. 4C,D). In one case where the retrained model performed better, it is able to recover all canonical WCF base pairs in a CRISPR RNA direct repeat (Fig. 4C), only missing a single U•G pair. However, the published SincFold model does not recover any pairs in the ground truth structure and proposes a non-canonical A•G pair. In a counterexample of a purine riboswitch, the retrained model fails to correctly predict any ground truth base pairs while the published model recovers all pairs except a non-canonical U•U pair (Fig. 4D). Taken together, these results highlight how the size of the training data and method for partitioning the sequences into rigorously separated training, validation, and test sets dramatically change how a given model architecture performs.

**Figure 4.**
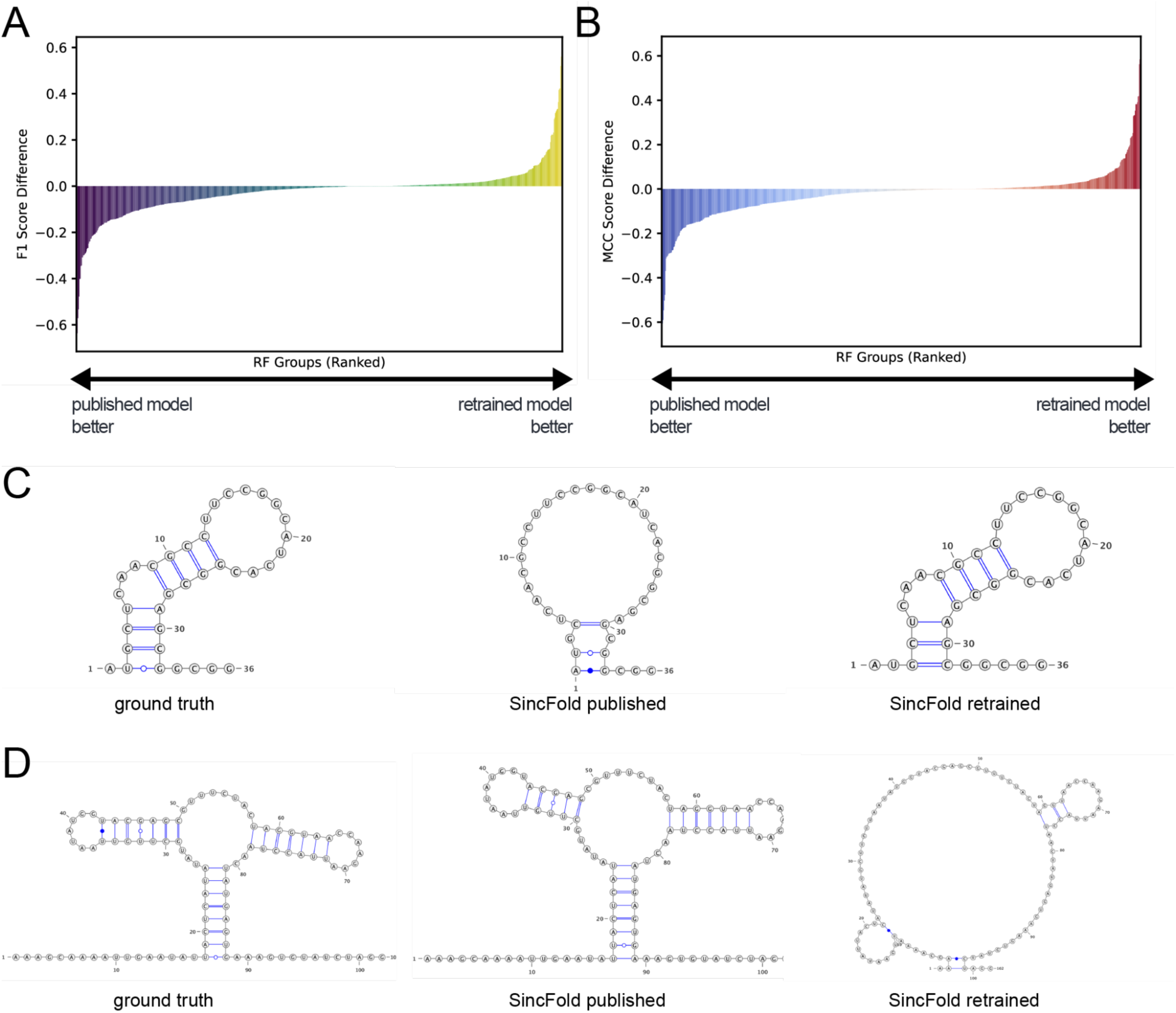
Differential performance across Rfam families by retrained SincFold. **A)** Rank order plot of per family average F1 score difference between the published SincFold model and the RNASSTR retrained SincFold model. Left-shifted families perform better with the published model and right-shifted families perform better with the RNASSTR retrieved model. **B)** Rank order plot of per family average MCC scores differences between published SincFold model and the RNASSTR retrained SincFold model. **C)** Representative RNA secondary structure of a family, RF01336 CRISPR RNA direct repeat, where the RNASSTR retrained model performed better than the published model. **D)** Representative RNA secondary structure of a family, RF00167 purine riboswitch, where the RNASSTR retrained model performed worse than the published model.

## Discussion

### Current Benchmark Datasets

Within the field of RNA secondary structure prediction, there currently exist three widely-used RNA secondary structure benchmark datasets: RNAStrAlign, ArchiveII, and bpRNA (41, 61, 71). Each of these has distinct characteristics in terms of size, sequence length distribution, and RNA family composition, and each has been utilized to various levels of success in training RNA ML models to predict RNA secondary structures.

RNAStrAlign (71) represents an alignment-based dataset aggregating known RNA secondary structures from 8 diverse RNA families (5S rRNA, tRNA, group I introns, 16S rRNA, tmRNA, SRP RNA, RNase P, and telomerase RNA) containing 37,149 sequence-structure pairs with lengths ranging from approximately 30 nucleotides to 1,851 nucleotides. While this represents a robust grouping of structurally diverse RNAs, this dataset only contains representative sequences from 8 structural families, limiting its utility in training general RNA structure models. Similarly, approximately 50% of the sequences in this dataset belong to 5S rRNA family, which limits the structural diversity despite the size of the dataset.

ArchiveII is a highly curated collection of 2,975 RNA sequences from 10 distinct RNA families (41) including several rRNAs as well as other common classes such as tRNA, RNase P, tmRNA, and self-splicing introns. While this dataset contains many fewer sequences than either RNAStrAlign (71) or bpRNA (61), it represents one of the older datasets for training and was originally compiled to provide a high-quality test set for RNA folding algorithms. Additionally, its curation ensures that each example is biologically relevant and non-redundant. For instance, ArchiveII contains representative rRNA sequences from different organisms rather than many near-duplicates. Because of its inclusion of large structured RNAs, ArchiveII is used as a stringent benchmark and has often been used as a hold-out test set in prior studies (i.e., models are sometimes trained on RNAStrAlign and evaluated on ArchiveII) (72, 73).

bpRNA represents the largest of the three datasets and the de facto standard for the field (61). This larger meta-database includes 102,318 RNA secondary structures, drawing from multiple sources, the largest being Rfam which contributes approximately half of the sequences and over 2,000 unique structural families. The scale of bpRNA makes it a popular choice for training deep learning models, but it carries an uneven family distribution. A few RNA families (tRNAs, 5S rRNAs) make up more than half of the sequences, while many other families are sparsely represented. To specifically test generalization to novel folds, an updated bpRNA-new dataset was introduced in Sato et al. (58) based on Rfam 14.2 (63). The bpRNA-new set contains sequences from approximately 1,500 new RNA families that were not present in the original bpRNA-1m compilation. By design, none of these families overlap with prior training sets, making bpRNA-new a benchmark for testing the cross-family generalization of current RNA secondary structure prediction models.

### The RNASSTR dataset

We mined three published RNA sequence databases to comprehensively guarantee diverse RNA fold representation in a new curated RNA secondary structure dataset we term RNASSTR. These databases, GTDB, NCBI RefSeq, and Rfam (Fig. 1) (63–65), represent an encompassing representation of our current understanding of biological sequence space. Using only reference genomes present in these data to limit the overrepresentation of model organisms, we identified 4,779,435 RNA sequences homologous to known RNA families defined in the Rfam database (Fig. 2). These sequences span all domains of life as well as viruses and represent a significantly larger sequence space than that queried in the three published datasets. Because previous datasets were limited in the folds they contained, we ensured that RNASSTR provides coverage of all RNA families present in Rfam, consisting of 4,028 unique folds representing all known RNA structural families. A notable feature of RNASSTR, much like bpRNA, is the overrepresentation of specific RNA families. As mentioned above, tRNAs account for 39.5% of the total sequences. However because of their short length this only accounts for 13.7% of the total nucleotides. Given training is performed on single nucleotide tokens we expect this overrepresentation to be mitigated, with the top 10 families by number of sequences making up a relatively more equal distribution of training families. In the future, it will be interesting to test alternative training schemes in which overrepresented families by number of sequences are subsampled, so family bias is less prevalent.

With RNASSTR, we are able to provide significantly more sequence depth and diversity compared to other existing RNA secondary structure datasets. A general trend in machine learning is improved model performance with data scaling (74). Because previous RNA secondary structure prediction models had been trained primarily with smaller and less diverse datasets, we hypothesized the lack of training data reduced their predictive power. However, upon retraining SincFoldusing RNASSTR we were not able to increase the average accuracy above that seen with the published model, suggesting there may be alternative aspects beyond the size of the training data driving lower model performance. However, we did observe a subset of families performed better with RNASSTR retrained models as compared to published models (Fig. 3,4). This suggests that the two versions of SincFold, the published version and the retrained version presented here, have learned different features in the data resulting in differential model performance. At this point, we have not been able to determine any specific features which may drive this difference (Fig. S3). Similarly, the performance differences between the SincFold models and RNAfold appear distinct, particularly for sequences where predictions failed to recover any correct base pairs, though further analysis of specific instances or features which allow thermodynamics-based models to outperform ML models, or vice versa represent an open question in the field.

One unsolved problem for both the published model parameters and the retrained model is data memorization. In deep learning, models can overfit the training data resulting in a pathology termed “memorization” (75), where instead of learning dataset features which allow the model to generalize, they instead memorize the sequence and secondary structures present in the training data. While we cannot directly comment on memorization in the published models, their large difference in performance on families on which they have been trained compared to novel families suggests model overfitting (42). Similarly, when retraining SincFold, the large discrepancy between the training and testing F1 and MCC scores (Table 1) also suggests this same pathology. One possible explanation for the difference in performance of the same model architecture trained on previous data versus RNASSTR could be the approach to splitting the data into training, test, and validation sets. In the case of secondary structure prediction, the approach used for bpRNA-1m, which performed data splitting using sequence identity rather than structural overlap, may artificially inflate F1 values in test sets. In other words, the RNASSTR test set used here to assess the published SincFold model may have some representation in the bpRNA-1m training data used to originally train SincFold.

To prevent a similar issue with RNASSTR, we performed a rigorous structure-based partitioning of the data into three sets, a training, a validation, and a testing set. By using the Rfam defined structural grammars, we ensured that these splits retained families which were mutually identifiable and remained in the same group. While the RNASSTR-retrained SincFold model did not outperform the published model parameters, the discrepancy between the training and testing F1 and MCC scores may represent a more accurate estimation of model generality compared to previous benchmarks. While the current RNA secondary structure prediction architectures we tested were unable to provide general RNA secondary structure prediction, we anticipate future models that make use of RNASSTR will be able to overcome this limitation and provide true generalization for RNA secondary structure prediction.

### Scaling Issues in current RNA secondary structure prediction models

Increased dataset size requires more efficient and faster training to make training models viable. Because RNASSTR is nearly 50x times larger than the most commonly used dataset, bpRNA-1m, the computational burden of training a single epoch using RNASSTR becomes much larger than with other datasets. The computational cost was highly limiting in the case of another model, MXFold2 (58) where each epoch required 268 GPU hours using a NVIDIA RTX A4500 GPU, and inference required 110 GPU hours on the same device. Training times of this length are infeasible, especially for academic groups with limited computational resources. As such we only completed 3 rounds of MXFold2 training before it became too resource-intensive to continue and we chose to devote more resources to the other model, SincFold. While SincFold was significantly quicker to train than MXFold2, each training epoch still required 64 GPU hours on a NVIDIA RTX A4500 GPU and 2 GPU hours for inference. A key aspect of improving future model architectures for RNA secondary structure prediction tasks would be to prioritize efficiency to allow for more throughput with the increasing sizes of training datasets. Models which enable parallel training using multiple GPUs should alleviate some of the burden of these large training sets where compute is not limiting.

RNASSTR provides a new step towards a standardized community wide benchmark for RNA secondary structure prediction, including rigorous methods for structure-based data splitting. However, it remains unclear whether sequence and secondary structure pairs will be sufficient to overcome the challenge of model generalization for RNA secondary structure prediction. For example, annotation of the full set of noncanonical base pairs in the secondary structure representations may be required. Furthermore, RNASSTR presently does not include information on pseudoknots, a key feature of many RNA structures that has been difficult to capture in training data and secondary structure prediction models. To date, since most secondary structures are determined computationally, it is not clear how to accomplish this extension. Alternatively, the incorporation of experimental data may improve the prediction accuracy of RNA secondary structure prediction models. RNA chemical probing data is scalable and high-throughout, with initial attempts to integrate it into RNA structure prediction pipelines already showing promise (51, 52).

## Data Availability

The code used to format the RNASSTR data, perform model retraining as well as inference scoring can be found at the following GitHub: https://github.com/romanagle/RNASSTR The RNASSTR dataset is hosted at Zenodo: https://doi.org/10.5281/zenodo.15319168

## Supplementary Data statement

A supplementary data file is provided with this manuscript.

## Acknowledgements

The authors would like to thank members of the Cate and Doudna Labs for their helpful discussions and insightful comments. The dataset filtering was inspired by similar analyses from Yekaterina Shulgina and would not have been possible without her suggestions.

## Author Contributions

J.H.C., J.A.D., and C.J.L conceptualized the project. C.J.L generated the RNASSTR dataset. C.J.L., T.K., R.N., C.M, and D.A.G. performed model retraining and assessment. C.J.L., T.K., R.N., C.M, and D.A.G. wrote the initial draft of the manuscript and all authors reviewed and edited the manuscript. J.H.C. and J.A.D. were responsible for funding and supervised the project.

---

**Supplementary Figure 1.**
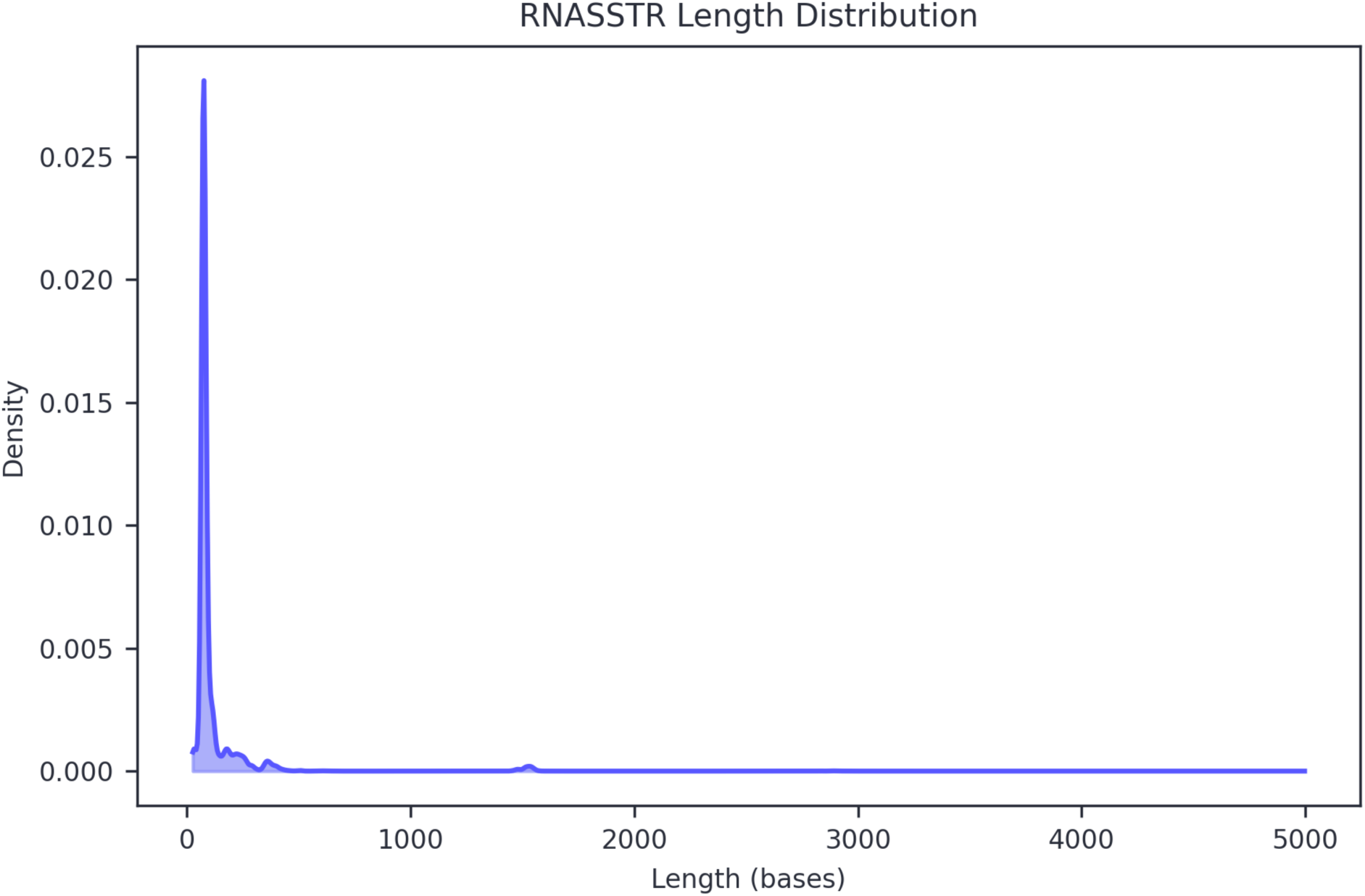
RNASSTR dataset sequence length distribution density plot. The peak at approximately 1550 nucleotides corresponds to bacterial small subunit rRNA. Longer sequences correspond to primarily bacterial and archeal large subunit rRNA.

**Supplementary Figure 2.**
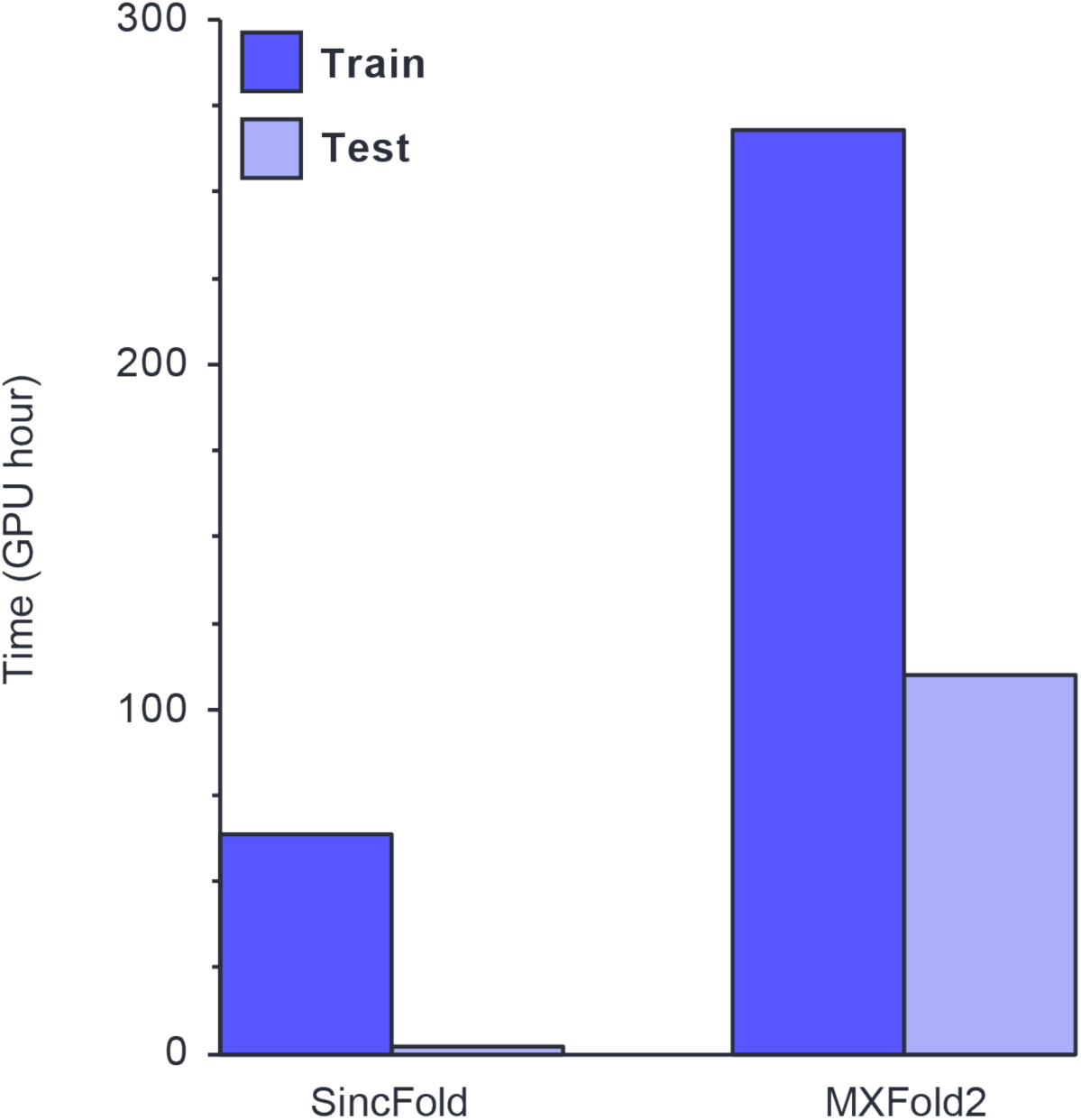
Model retraining times. SincFold (left) and MXFold (right) training and testing times per epoch in GPU hours using a single NVIDIA RTX A4500 GPU.

**Supplementary Figure 3.**
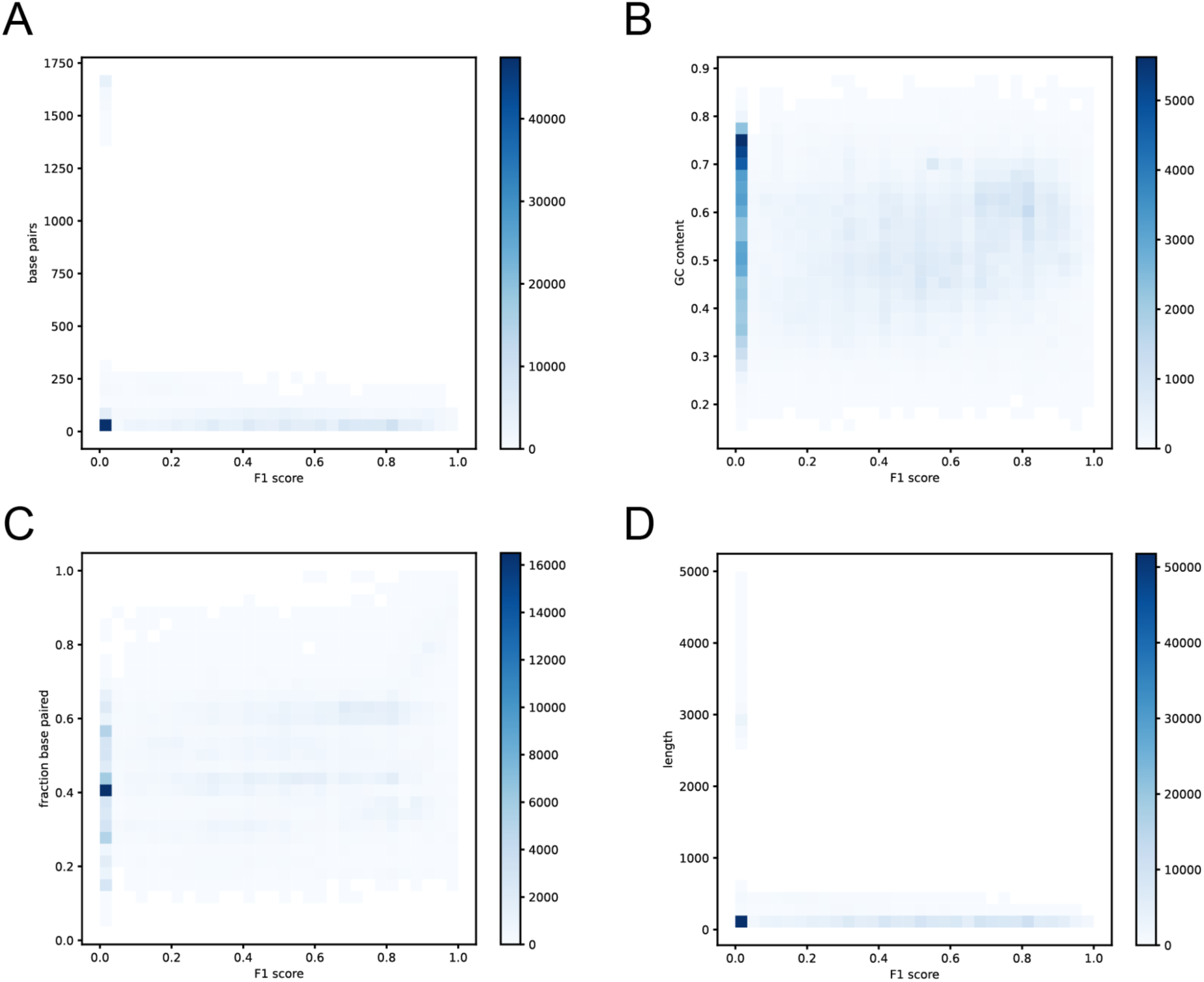
RNASSTR features versus F1 score. **A-D)** Density of RNASSTR test partition features versus the resulting F1 score from the retrained SincFold model. Features analyzed are as follows: absolute number of base pairs in the ground truth structure (**A**), sequence GC content (**B**), fraction of bases paired in the ground truth structure (**C**), and sequence length (**D**).

**Supplementary Figure 4.**
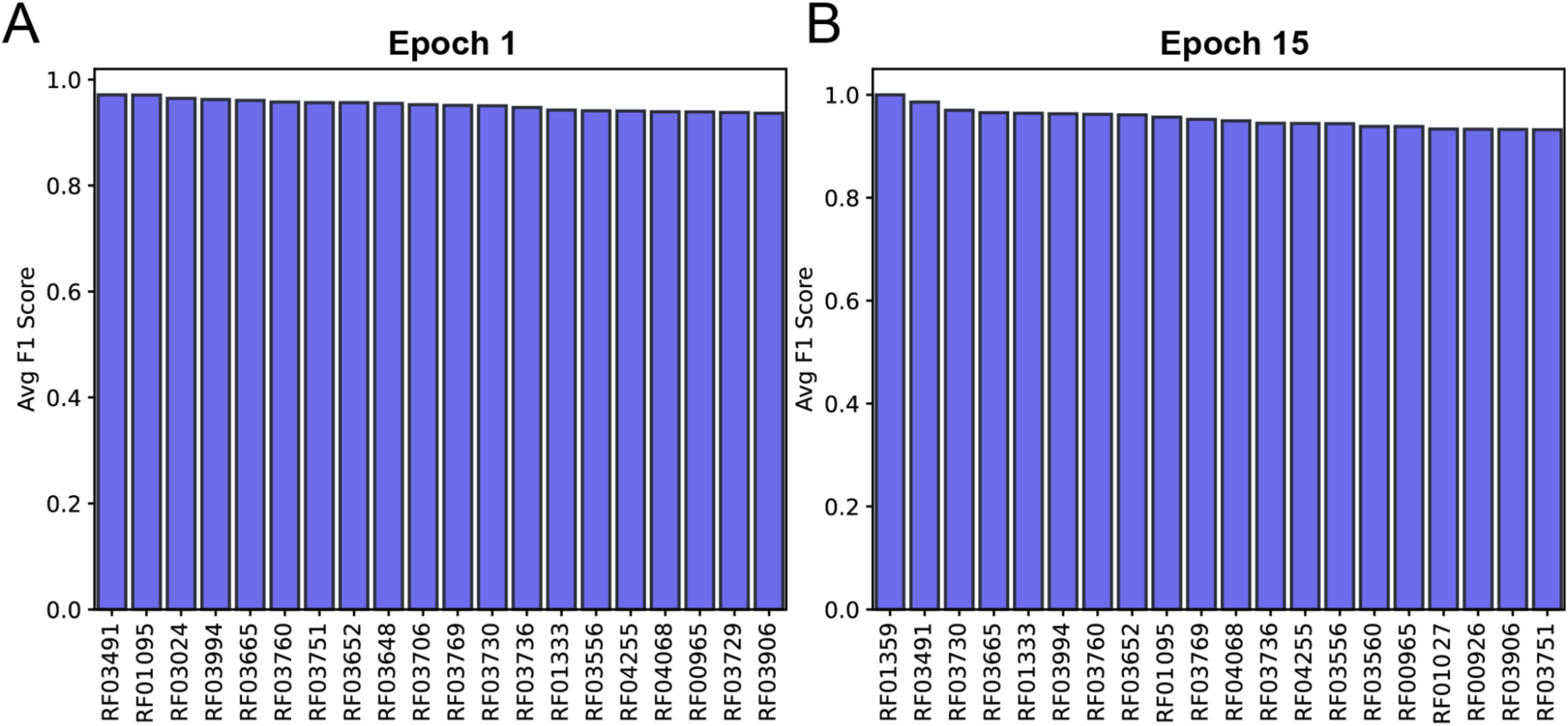
Average F1 score of top 20 performing classes. Shown are the F1 scores predicted by RNASSTR retrained SincFold at the end of epoch 1 (**A**) and epoch 15 (**B**).

**Supplementary Figure 5.**
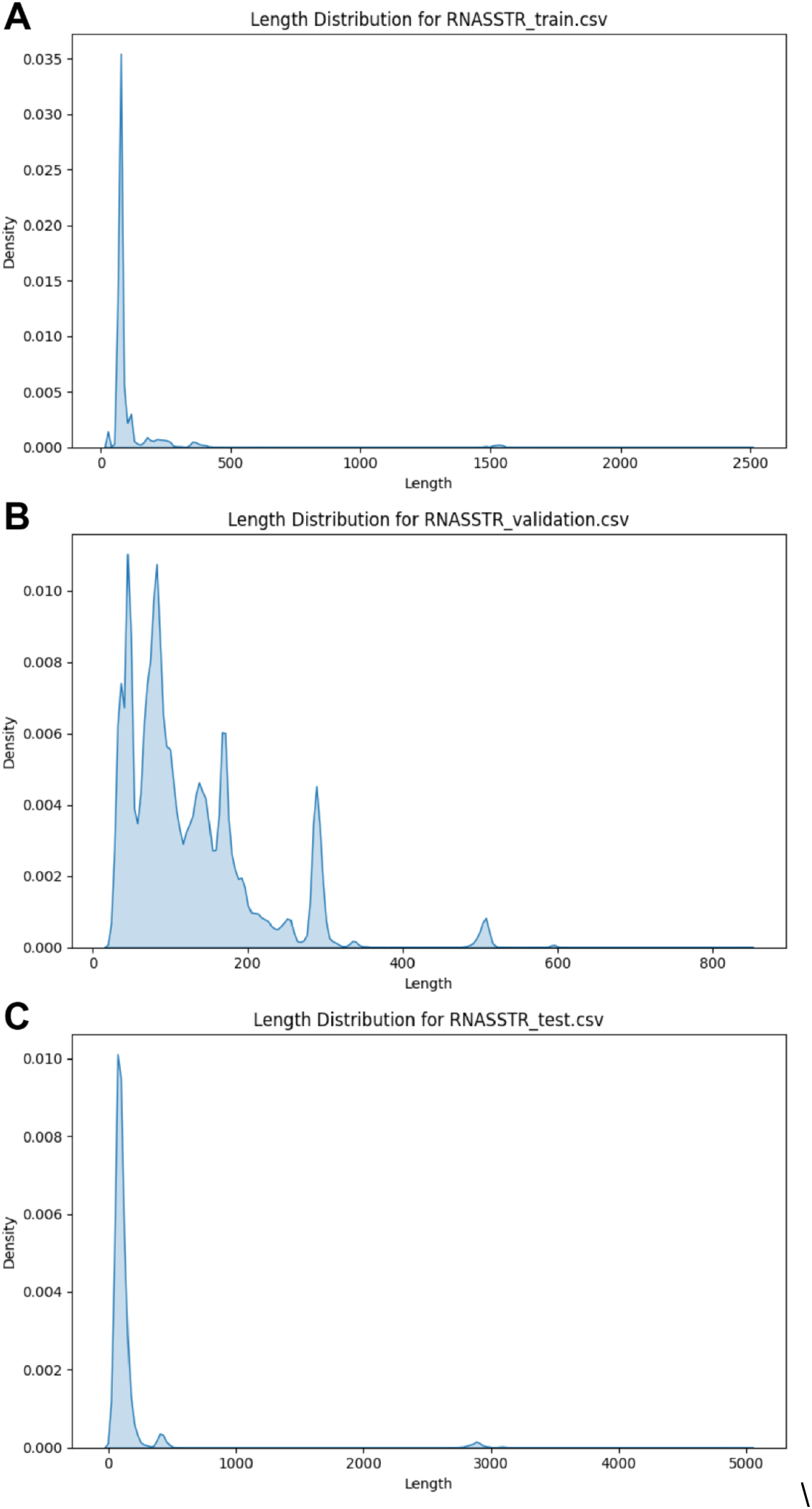
RNASSTR features versus F1 score. **A-C)** Length distributions of the sequences contained in the three data partitions of RNASSTR: train (**A**), validation (**B**), and test (**C**). Note that the length scales on the horizontal axis differ for each partition, based on the longest sequence present.

